# Identification of Bone Morphogenetic Protein 7 as a Master Regulator of Gastric Cancer-Associated Cachexia

**DOI:** 10.64898/2026.04.13.717813

**Authors:** Mayu Yasuda-Koiwa, Takahiro Shoda, Akiho Nishimura, Tadahito Yasuda, Atsuko Yonemura, Kota Muraki, Yuya Okamoto, Takuya Tajiri, Y Alan Wang, Takatsugu Ishimoto

## Abstract

Cachexia is a devastating and multifactorial syndrome characterized by progressive loss of body weight, skeletal muscle wasting, and systemic inflammation, frequently observed in patients with advanced gastric cancer (GC) with peritoneal dissemination. Despite its clinical significance, the molecular mechanisms underlying cancer-associated cachexia remain poorly understood. In this study, comparative transcriptomic analysis using the GEMINI database identified bone morphogenetic protein 7 (BMP7) as a novel candidate cachexia-inducing factor, along with the known cachexia mediators, growth differentiation factor 11 (GDF11) and growth differentiation factor 15 (GDF15). Functional studies demonstrated that BMP7 acts as an upstream regulator that drives cachectic phenotypes by inducing the expression of GDF11 and GDF15. Knockdown of BMP7, GDF11, or GDF15 in the cachexia-inducing GC cell line, MKN45 significantly attenuated weight loss and muscle wasting *in vivo*. Conversely, overexpression of BMP7 in the non-cachectic GC cell line, NUGC3 induced cachexia and upregulated GDF11 and GDF15 in tumor tissues. Furthermore, clinical analysis revealed that high BMP7 expression in tumor specimens from patients with advanced GC was associated with significantly poorer overall survival. These findings identify BMP7 as a master regulator of cancer-associated cachexia through the induction of GDF11 and GDF15 and suggest its potential as a promising therapeutic target for mitigating cachexia in GC.

## Introduction

Cachexia is a devastating, multifactorial, and often irreversible systemic condition characterized by weight loss, wasting of muscle and adipose tissue wasting, and systemic inflammation that is often associated with anorexia and increased energy expenditure [1]. This syndrome directly affects patients’ quality of life and limits both the efficacy and availability of cancer therapies. Cachexia is most commonly observed in patients with advanced-stage cancer, affecting up to 80% of this population, and is a major contributor to cancer-related morbidity and mortality [2]. Despite extensive research efforts, no effective therapies have been established, largely due to an incomplete understanding of the underlying molecular mechanisms.

The incidence of cachexia varies by tumor type, occurring in over 80% of patients with gastric or pancreatic cancer [3]. One contributing factor is that gastric cancer (GC), particularly in its advanced stages, frequently leads to peritoneal dissemination, which is often accompanied by systemic inflammation [4–6]. This inflammatory response plays a key role in driving the cachectic phenotype, characterized by severe weight loss, muscle wasting, and anorexia. These clinical features suggest that tumor-secreted factors—particularly those present in the peritoneal microenvironment—play a critical role in the pathogenesis of cachexia. Among such factors, members of the transforming growth factor-β (TGF-β) superfamily—including growth differentiation factor 11 (GDF11) and growth differentiation factor 15 (GDF15)—have been reported to mediate systemic wasting and to be associated with cachexia [7, 8]. GDF11 directly contributes to skeletal muscle wasting by activating phosphorylated SMAD2/3, leading to significant atrophy of differentiated myotubes [9, 10]. In contrast, GDF15 binds to its specific receptor, glial cell line-derived neurotrophic factor (GDNF) family receptor alpha-like (GFRAL), in the hindbrain, resulting in reduced food intake and weight loss [11–13]. Especially for GDF15, ongoing clinical trials are investigating the therapeutic potential of targeting the GDF15-GFRAL signaling axis [14]. While several tumor-derived molecules are believed to contribute to this process, the precise molecular drivers remain poorly understood.

The present study was conducted to identify a master regulator of cancer-associated cachexia. Our findings demonstrate that bone morphogenetic protein 7 (BMP7), a members of the TGF-β superfamily, act as an upstream regulator by inducing the expression of GDF11 and GDF15, two key mediators of the cachectic phenotype. This discovery is further supported by our observation that high BMP7 expression correlates with increased cachexia and poorer survival, suggesting it may represent a promising therapeutic target for future interventions.

## Results

### MKN45 possesses particular mechanisms inducing peritoneal engraftment and the development of cachexia

We first generated a peritoneal dissemination mouse model using diffuse-type GC cell lines: OCUM12, KATOIII, NUGC3, OCUM-2MD3, and MKN45. As reported previously, MKN45 is known to form robust peritoneal metastases and induce cachexia in the nude rat model [15]. Over the 30-day observation period, mice bearing NUGC3, OCUM-2MD3, and MKN45 developed intraperitoneal tumors (Figure S1A). However, only the MKN45-bearing mice exhibited significant body weight loss, and a markedly reduced survival rate compared to mice bearing other diffuse-type GC cell lines (Figure 1A and Figure S1B). To determine whether this body weight loss met the criteria for cachexia, we compared MKN45-bearing mice with OCUM-2MD3-bearing mice, which served as a non-cachectic controls. Notably, the quadriceps muscle mass of MKN45-bearing mice was significantly reduced at both the macroscopic and microscopic levels (Figures 1B and 1C). In addition, grip strength was significantly decreased in MKN45-bearing mice relative to the OCUM-2MD3 bearing mice (Figure 1D). To further validate these findings at the molecular level, we examined additional muscle atrophy markers in the gastrocnemius and tibialis anterior muscles. MKN45-bearing mice showed significantly increased mRNA expression of *MuRF1* and *Atrogin-1* compared with OCUM-2MD3-bearing mice in both muscle groups (Figure 1E). Moreover, to evaluate whether reduced food intake contributed to body weight loss, we measured the food intake. MKN45-bearing mice showed a significant reduction in food intake compared with OCUM-2MD3-bearing mice (Figure 1F). Furthermore, to assess changes in adipose tissue, we quantified white adipose tissue (WAT), which primarily serves as an energy-storing depot. The inguinal WAT index (percentage of inguinal WAT weight relative to total body weight) was markedly reduced in MKN45-bearing mice compared with OCUM-2MD3-bearing mice. In addition, the inguinal WAT exhibited a reddish appearance consistent with browning, a process in which WAT acquires brown adipose tissue (BAT)-like characteristics, a tissue specialized for thermogenic energy expenditure (Figure 1G and 1H). These results indicate that only MKN45-bearing mice developed cachexia secondary to peritoneal metastasis.

**Figure 1.**
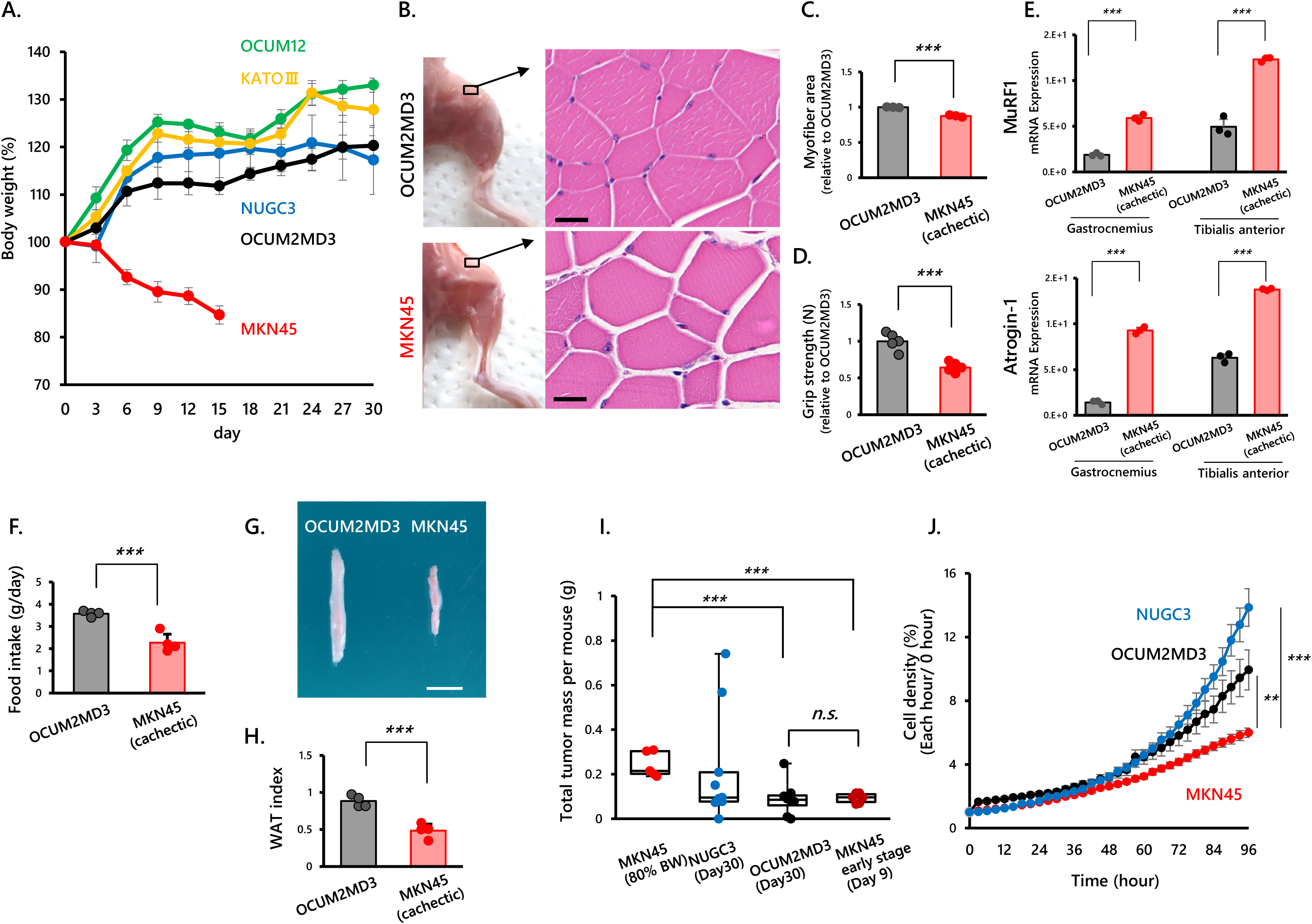
MKN45 cells display specific capabilities that drive peritoneal engraftment and cachexia. A. Body weight changes over 30 days following intraperitoneal injection of diffuse-type GC cell lines (n=4 mice per group). In the MKN45 group, monitoring was discontinued before day 30 because body weight fell below 80% of the initial value. B. Representative images of quadriceps muscle from mice bearing MKN45 or OCUM-2MD3 tumors at the time of sacrifice. Muscles were harvested at the time of sacrifice, which was performed when predefined humane endpoints were reached. Scale bars, 20 μm C. Quantification of myofiber area from the representative images in (B). D. Grip strength measurements in each group measured at the time of sacrifice (OCUM-2MD3: n = 5 mice; MKN45: n = 6 mice). E. qRT-PCR analysis of muscle atrophy–related genes (*MuRF1* and *Atrogin-1*) in the gastrocnemius and tibialis anterior muscles of each group (OCUM-2MD3: n = 3; MKN45: n = 3). F. Food intake measurements in each group at day between day 14 and day 15 after tumor implantation (n=4 mice per group). G. Representative images of inguinal WAT from mice bearing MKN45 or OCUM-2MD3 tumors at the time of sacrifice. Scale bars, 10 mm. H. Quantification of the inguinal WAT fat index (percentage of the combined bilateral inguinal WAT weight relative to total body weight) in mice bearing MKN45 or OCUM-2MD3 tumors at the time of sacrifice (OCUM-2MD3: n = 4 mice; MKN45: n = 4 mice). I. Quantification of tumor mass in each group measured at the time of sacrifice (MKN45: n=5 mice; NUGC3: n=9 mice; OCUM-2MD3: n = 8 mice; MKN45 early stage: n = 6 mice). J. In vitro growth assay of diffuse-type GC cell lines over 96 hours. n.s., not significant; **p < 0.01; ***p < 0.001; Mann–Whitney U test.

Subsequently, we compared the tumor mass at the time of sacrifice in mice bearing NUGC3, OCUM-2MD3, and MKN45. Tumor mass tended to be greater in MKN45 bearing mice than in OCUM-2MD3-bearing mice. To clarify that cachexia induction in our model is not solely attributable to tumor burden, we additionally evaluated MKN45-bearing mice at an earlier time point, 9 days after MKN45 cell transplantation. Although differences in body weight were already evident at this stage (Figure 1A), early-stage MKN45-bearing mice (day 9) did not exhibit a significant increase in tumor mass compared with OCUM-2MD3-bearing mice (Figure 1I). These results indicate that body weight loss occurs even when the tumor size remains small, supporting the notion that weight loss in MKN45-bearing mice cannot be explained solely by tumor mass. Moreover, our *in vitro* analyses showed that MKN45 cells proliferate more slowly than other diffuse-type gastric cancer cell lines (Figure 1J), further supporting the independence of the cachexia phenotype from tumor growth rate. Collectively, these findings suggest that MKN45 exerts unique effects by secreting humoral factors that promote peritoneal engraftment and trigger the onset of cachexia.

### Candidate genes potentially involved in cachexia induction are identified via GEMINI Database

To identify humoral factors secreted by GC cells that may contribute to cachexia, we utilized the GEMINI database (GSE22183), which contains genome-wide mRNA expression profiles for 37 unique GC cell lines established by Tan et al. [16]. Using this microarray dataset, we performed a comparative gene expression analysis between the cachexia-inducing cell line MKN45 and two non-cachexia-inducing cell lines, NUGC3 and KATOIII. This comparative analysis effectively allowed us to evaluate differences in cytokine- and chemokine-related gene expression among distinct GC cell types. This analysis generated 13 candidate genes potentially associated with cachexia (Figure 2A). We then validated the expression of these 13 candidate genes in five GC cell lines using RT-PCR. Among them, we identified five genes (*BMP7*, *GDF15*, *GDF11*, *THNSL2*, and *TNFSF13*) as strong candidates for cachexia inducers. These genes were consistently and significantly upregulated in MKN45 compared to the other cell lines, with their expression levels maintained at comparably high levels (Figure 2B).

**Figure 2.**
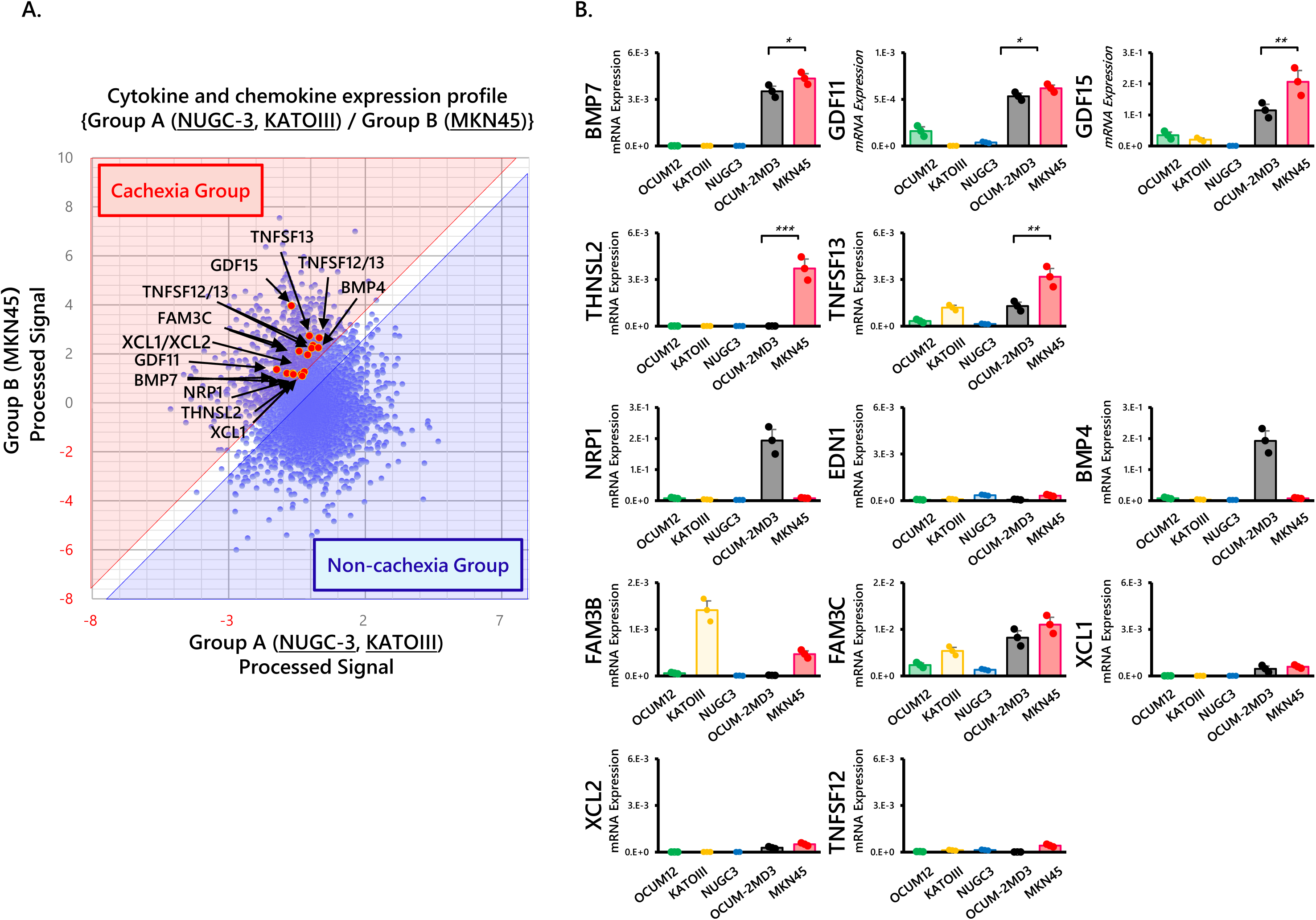
Candidate genes implicated in cachexia induction were identified using the GEMINI database. A. Cytokine expression profile comparing group B (MKN45) with group A (NUGC3 and KATOIII) using the GEMINI database (GSE22183). Red points indicate cytokines; blue points indicate other genes. B. Expression of selected candidate genes validated by qRT-PCR. *p < 0.05; **p < 0.01; ***p < 0.001; Mann–Whitney U test.

### *BMP7* induces cachexia by upregulating *GDF11* and *GDF15* expression

To investigate the roles of the candidate genes in the development of cachexia, we established stable MKN45 cell lines with shRNA-mediated knockdown of *BMP7*, *GDF11*, *GDF15*, *THNSL2*, and *TNFSF13* (Figure S2A), and performed *in vivo* experiments using mice transplanted with these knockdown cells. Mice bearing sh*THNSL2* and sh*TNFSF13* cells exhibited cachectic phenotypes comparable to those observed in the sh-control group (data not shown). In contrast, knockdown of *BMP7*, *GDF11*, or *GDF15* resulted in attenuation of cachexia, as evidenced by preserved body weight and grip strength (Figures 3A, 3B and 3C). To further characterize the effects of BMP7 knockdown on skeletal muscle, we performed additional analyses focusing on shBMP7 mice. Quantification of quadriceps muscle mass demonstrated a significant restoration in shBMP7 mice compared with the sh-control group (Figure 3D).

**Figure 3.**
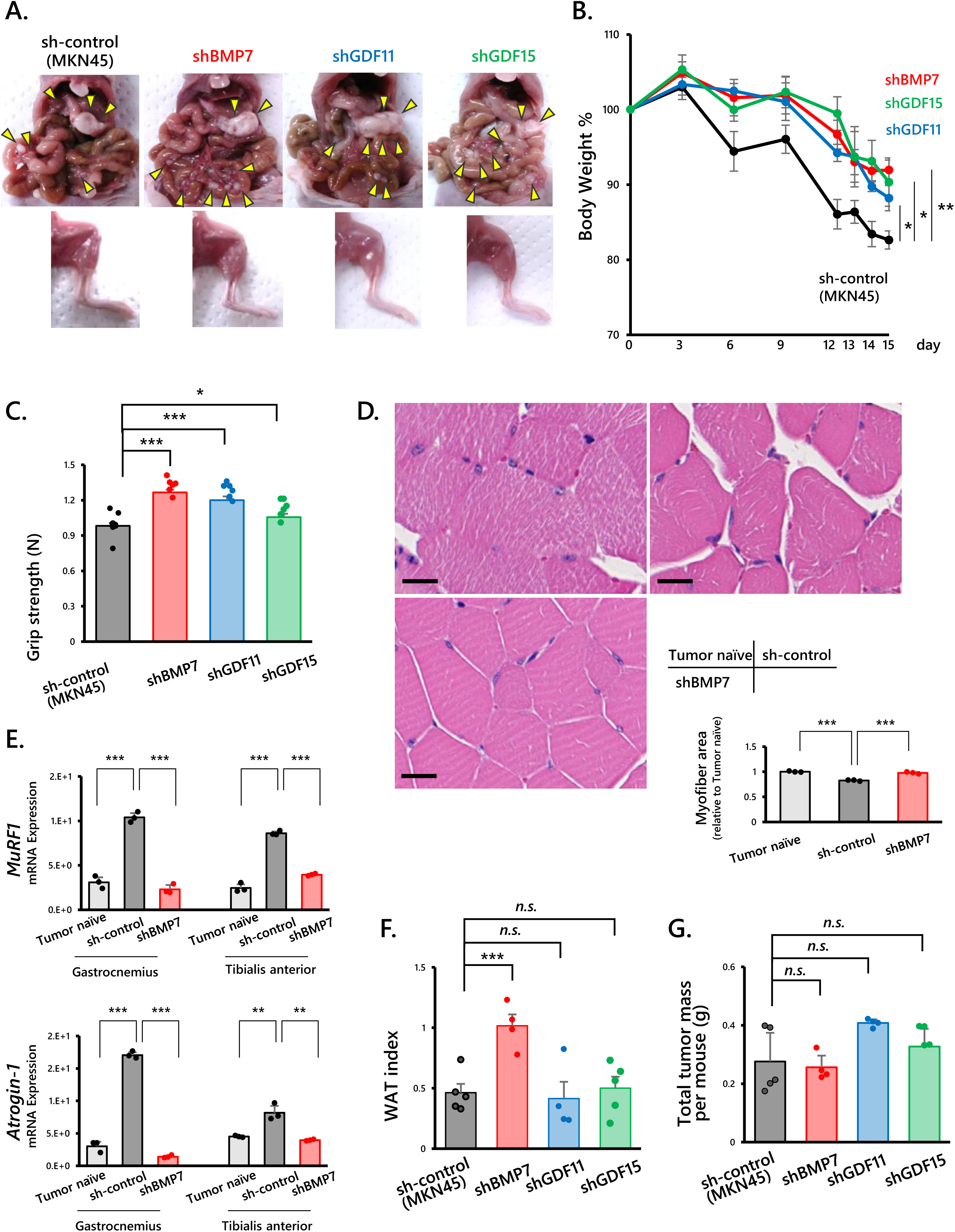
Knockdown of BMP7, GDF11, or GDF15 attenuates cachexia-associated phenotypes without detectable changes in tumor burden. A. Representative images of tumor-bearing mice and quadriceps muscle after intraperitoneal injection of MKN45 cells with shRNA-mediated knockdown of BMP7, GDF11, GDF15, or control shRNA (shBMP7: n=4 mice; shGDF11: n=5 mice; shGDF15: n=6 mice; sh-control: n=6 mice). B. Body weight changes over 15 days in each group. C. Grip strength measurements in each group (n=6 mice per group). D. Representative images of quadriceps muscle in each group (Tumor naïve, sh-control, shBMP7). Muscles were harvested at the time of sacrifice. Scale bars, 20 μm. The accompanying graph shows quantification of myofiber area. E. qRT-PCR analysis of muscle atrophy–related genes (MuRF1 and Atrogin-1) in the gastrocnemius and tibialis anterior muscles of each group (Tumor naïve: n = 3; sh-control: n = 3; shBMP7: n=3). F. Quantification of the inguinal WAT fat index (percentage of the combined bilateral inguinal WAT weight relative to total body weight) in each group (shBMP7: n=4 mice; shGDF11: n=4 mice; shGDF15: n=5 mice; sh-control: n=5 mice). G. Quantification of tumor mass in each group (shBMP7: n=4 mice; shGDF11: n=4 mice; shGDF15: n=4 mice; sh-control: n=5 mice). n.s., not significant; *p < 0.05; **p < 0.01; ***p < 0.001; Mann–Whitney U test.

Consistently, in the gastrocnemius and tibialis anterior muscles, shBMP7 mice exhibited significantly reduced mRNA expression of the muscle atrophy markers *MuRF1* and *Atrogin-1* compared with sh-control mice (Figure 3E). Regarding the changes in adipose tissue, inguinal WAT index was substantially restored in shBMP7 mice compared with the sh-control group, whereas no appreciable improvement was observed in shGDF11 or shGDF15 mice (Figure 3F). These results indicate that BMP7 plays a predominant and direct role in regulating adipose tissue remodeling in this model, whereas the BMP7– GDF11/GDF15 axis is not a major determinant of WAT mass. Moreover, tumor volumes did not differ significantly between the shBMP7 and sh-control groups, or between the shGDF11, shGDF15, and their respective control groups (Figure 3G). This similarity in tumor burden is consistent with the notion that these knockdowns do not substantially affect tumor growth and supports the conclusion that the observed cachexia is independent of tumor mass.

To further investigate the regulatory relationships among these three genes, we examined the expression levels of *GDF11* and *GDF15* in sh*BMP7* cells. Both genes showed reduced expression compared to sh-control cells, and this suppression was rescued by treatment with recombinant BMP7, suggesting that BMP7 positively regulates GDF11 and GDF15 expression (Figure 4A). In contrast, knockdown of *GDF11* or *GDF15* did not affect *BMP7* expression levels, indicating that GDF11 and GDF15 are downstream targets of BMP7 (Figure 4B). To further support a mechanistic link between BMP7 signaling and GDF11/GDF15 regulation, we next examined activation of the SMAD pathway, a canonical downstream pathway of BMP signaling, by Western blot analysis. Phosphorylation level of SMAD1/5 was significantly reduced in sh*BMP7* cells compared with sh-control cells, and this reduction was restored by recombinant BMP7 treatment (Figure 4C).

**Figure 4.**
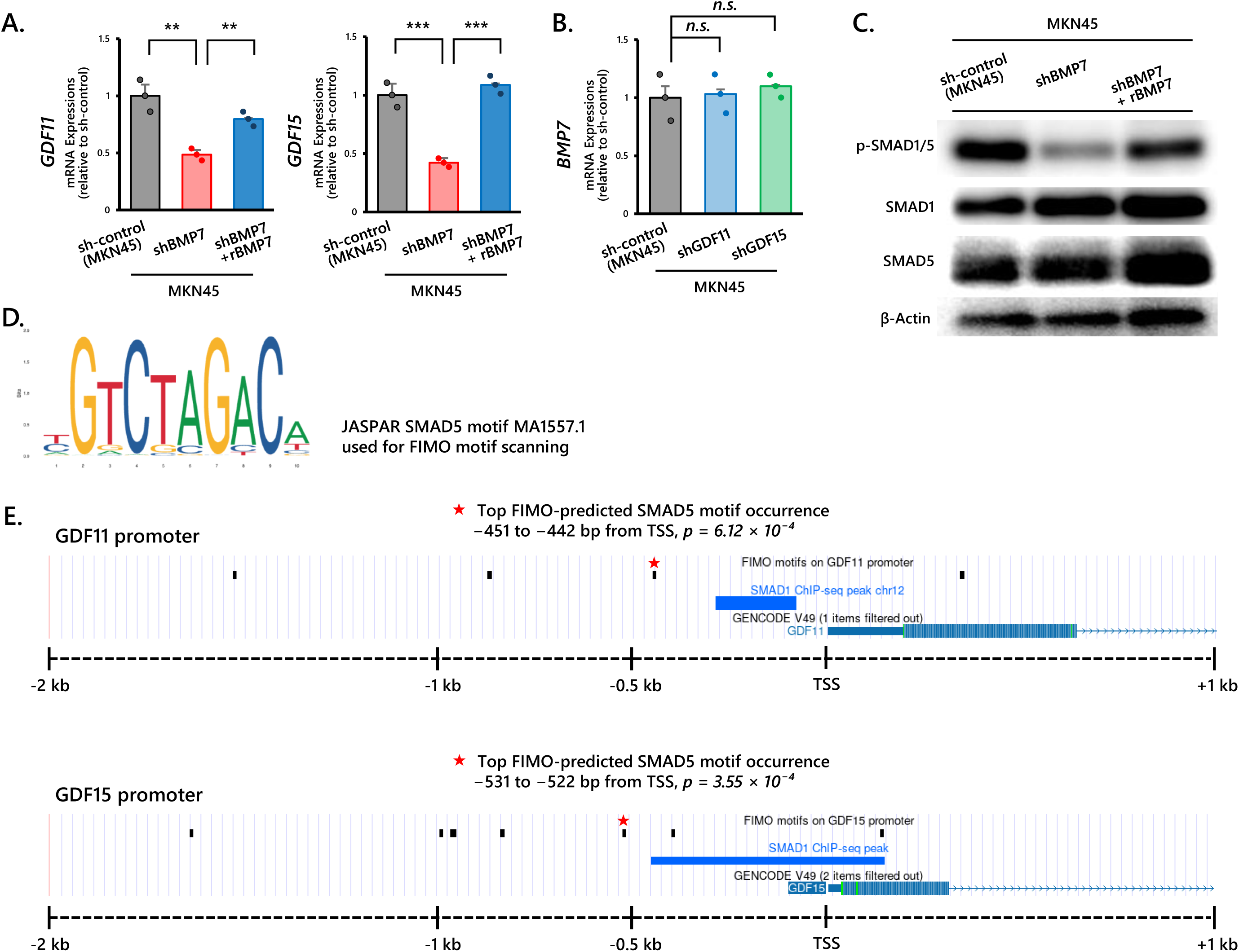
BMP7-dependent regulation of GDF11 and GDF15 expression and identification of candidate SMAD5-binding motifs in their promoters. A. Expression of GDF11 and GDF15 analyzed by qRT-PCR in sh-control, shBMP7, and shBMP7 + recombinant BMP7 (rBMP7) MKN45 cells. B. BMP7 expression analyzed by qRT-PCR in sh-control, shGDF11, and shGDF15 MKN45 cells. C. Western blot analysis of pSMAD1/5 level in shBMP7 cells with or without treatment with rBMP7. Total SMAD1/5 levels were analyzed as loading controls. D. Sequence logo of the JASPAR SMAD5 motif (MA1557.1) used for FIMO motif scanning. E. In silico analysis of the GDF11 and GDF15 promoter regions (−2 kb to +1 kb relative to the TSS). Black ticks indicate predicted SMAD5 motif occurrences predicted by FIMO. Red stars the top-ranked motif occurrence in each promoter. Because a comparable public SMAD5 ChIP-seq dataset was not identified, publicly available SMAD1 ChIP-seq peaks are shown for positional comparison. n.s., not significant; **p < 0.01; ***p < 0.001; Mann–Whitney U test.

**Figure 5.**
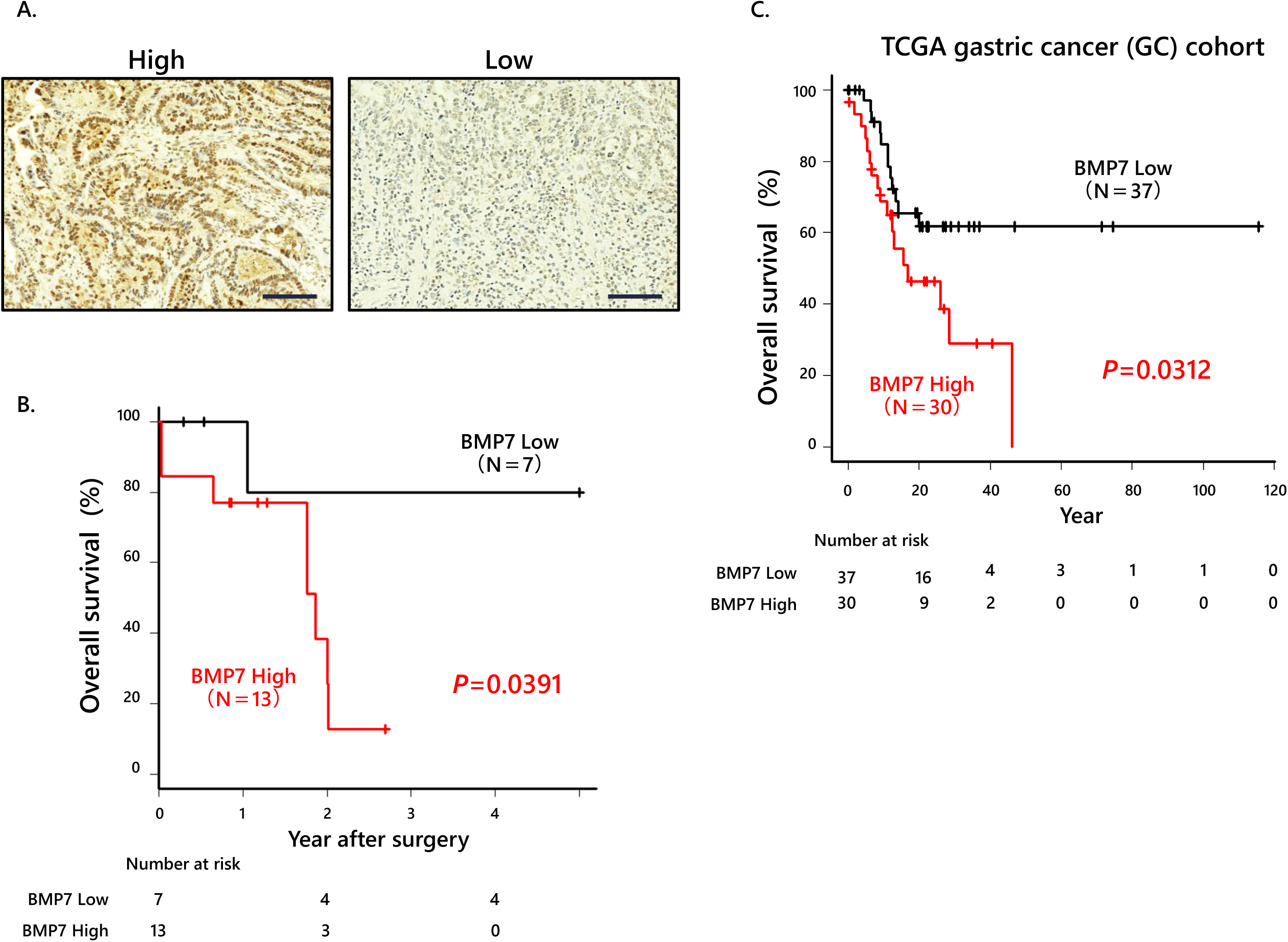
BMP7 promotes tumor malignancy in advanced gastric cancer patients with cachexia. A. Representative IHC images of BMP7 staining in tumor samples from advanced GC patients. Scale bars, 100 μm. B. Overall survival of 20 advanced GC patients stratified by BMP7 expression. C. Overall survival of advanced GC patients from TCGA database stratified by BMP7 mRNA expression levels (high, n=30; low, n=37).

We next examined whether the GDF11 and GDF15 promoter regions contain putative SMAD-responsive elements. Using the JASPAR SMAD5 motif (Figure 4D), *in silico* motif analysis identified candidate SMAD5-binding motifs within the promoter regions of both genes. The highest-scoring motifs were located at −451 to −442 bp upstream of the transcription start site (TSS) in the GDF11 promoter and −531 to −522 bp upstream of the TSS in the GDF15 promoter. Publicly available SMAD1 ChIP-seq peaks were located near the predicted SMAD5 motif sites in both promoters (Figure 4E). Although these analyses do not demonstrate direct SMAD1/5 binding, they support the presence of putative SMAD-responsive regulatory regions within the GDF11 and GDF15 promoters. Together, these findings suggest that BMP7-dependent activation of SMAD signaling is associated with the induction of GDF11 and GDF15 expression, consistent with the hypothesis that BMP7 functions as an upstream regulator of cachexia-associated signaling in this model.

### BMP7 promotes tumor malignancy in advanced GC patients with cachexia

To determine the clinical relevance of our findings in advanced GC patients with cachexia, we evaluated the association between BMP7 expression levels in tumor tissues and patient outcomes, including 5-year overall survival (OS) using Kaplan–Meier analysis. We first examined BMP7 expression in tumor samples from 20 advanced GC patients. Patients with high BMP7 expression in tumor areas exhibited significantly poorer OS compared to those with low BMP7 expression (Figure 4A and 4B, Table S1).

To further validate these observations in an independent cohort, we analyzed publicly available data from The Cancer Genome Atlas (TCGA). Consistent with our institutional cohort, higher BMP7 mRNA expression was significantly associated with worse OS in patients with diffuse-type advanced GC in the TCGA cohort (Figure 4C). Together, these findings indicate that elevated BMP7 expression is associated with an aggressive disease course and poor prognosis in advanced GC, supporting the clinical relevance and generalizability of our experimental observations.

## Discussion

BMP7, also known as osteogenic protein-1 (OP-1), is a secreted cytokine belonging to the TGF-β superfamily. It was originally identified as a factor inducing ectopic chondro-osteogenesis and plays a key role in skeletal development, bone remodeling, and fracture repair [17]. Recombinant human BMP7 has therefore been used therapeutically for bone fractures [18]. However, the functions of BMP7 extend beyond skeletal tissues, contributing to embryonic development and systemic homeostasis. Mice deficient in BMP7 exhibit profound developmental abnormalities across multiple organs [19–21]. Moreover, BMP7 is involved in appetite regulation. BMP7 has functions as an anorexigenic factor via a leptin-independent, which is central mTOR pathway in the hypothalamus to prevent obesity [22].

While BMP7 is essential for survival and physiological homeostasis, the biological effects of TGF-β family proteins are highly context-dependent. BMP7 expression has been reported across multiple malignancies and is frequently associated with metastasis and poor prognosis in cancer patients [23, 24]. In GC, BMP7 has also been implicated in tumor progression [25]. Moreover, BMP7 secreted by tumor cells acts on macrophages and CD4⁺ T cells within the tumor microenvironment, suppressing MAPK14 expression and impairing proinflammatory immune responses. Notably, BMP7 blockade in combination with anti–PD-1 therapy has demonstrated synergistic antitumor effects in *in vivo* models [26], highlighting the critical role of interactions between the tumor and immune microenvironment. Given that our demonstration in this study that the BMP7 secreted from GC cells induces the cachectic condition, BMP7 inhibition in advanced-stage GC may ameliorate cachectic symptoms by modulating appetite and energy homeostasis through regulating the GDF11 and GDF15 signaling. Although our data show that BMP7 activates SMAD1/5 signaling and increases GDF11 and GDF15 expression, the transcriptional link between these events remains to be fully defined. *In silico* analyses identified candidate SMAD5 motif occurrences and nearby public SMAD1 ChIP-seq peaks in both promoters, consistent with the presence of putative SMAD-responsive regulatory regions. However, these findings do not establish direct SMAD1/5 binding or promoter activation in our experimental system. Thus, whether BMP7 regulates GDF11 and GDF15 transcription directly or indirectly through intermediate regulatory factors remains to be clarified. Importantly, GDF15 is a secreted protein currently under clinical investigation as a potential therapeutic target for cachexia [27, 28]. In this context, our findings provide mechanistic insight into the role of BMP7 in cancer-associated cachexia and highlight its translational relevance.

Another important function of certain BMP family members, including BMP7, has also been shown to regulate adipose tissue homeostasis. BMP7 plays a key role in the differentiation and maintenance of adipocytes, particularly BAT [29]. BAT is specialized for energy expenditure and thermogenesis, releasing energy as heat via uncoupled mitochondria. Activation of thermogenesis in the interscapular BAT has been observed in a syngeneic mouse tumor transplant model and suggested to contribute to the hypermetabolic state of cachexia [30]. On the other hand, WAT primarily serves as an energy storage depot, accumulating triglycerides in intracellular lipid droplets and providing them through lipolysis when energy is needed. WAT and BAT usually perform opposite physiological functions; WAT promotes energy storage, while BAT facilitates energy expenditure through thermogenesis. Moreover, a phenotypic switch from WAT to BAT, known as WAT browning, has been reported during the initial stages of cancer-associated cachexia, as demonstrated in the present study. This process precedes skeletal muscle atrophy and is characterized by increased expression of uncoupling protein 1 (UCP1), which shifts mitochondrial respiration from ATP synthesis to heat production [31].

In addition to the role of BMP7, another perspective worth considering is the function of bone morphogenetic protein 4 (BMP4) in regulating adipose tissue homeostasis. BMP4, a member of the TGF-β superfamily, has been shown to influence adipocyte development by promoting the conversion of brown pre-adipocytes into cells with white fat-like characteristics [32]. Moreover, transgenic overexpression of BMP4 specifically in WAT results in decreased WAT mass and smaller white adipocytes, alongside an increased population of adipocytes exhibiting molecular and morphological features of BAT [33]. These suggest that BMP4 may play a role in modulating the balance between energy-storing and energy-expenditure adipocyte populations, potentially exerting effects that are opposite to those of BMP7. This contrasting role between BMP4 and BMP7 may have important implications for the regulation of systemic energy homeostasis in the context of cancer-associated cachexia.

In our study, we also observed that BMP7 expression in OCUM-2MD3 cells is relatively high compared to other GC cell lines, as well as MKN45 cells. While OCUM-2MD3 cells successfully engrafted in host mice, they did not induce a cachectic phenotype. Interestingly, BMP4 expression was significantly elevated in OCUM-2MD3 cells (Figure 2B), raising important questions about the potential interplay between BMP7 and BMP4. How the disruption of the balance between BMP7 and BMP4 influences cachexia progression and tumor development remains unclear. This warrants further investigation as a key area of future research. Taken together, these findings support the notion that cancer-associated cachexia is unlikely to be driven by a single downstream effector. Instead, there is an urgent need to identify a further upstream regulatory molecule, a potential “master switch” that coordinates the cachectic program. Uncovering such a mechanism would provide critical insight and could offer a novel therapeutic target to improve clinical outcomes in cachectic patients with GC.

A limitation of this study is that all animal experiments were conducted using male mice. Several recent reports have demonstrated that cancer cachexia exhibits sex-dependent differences in its prevalence, severity, metabolic adaptations, and patterns of muscle wasting in both preclinical models and patients [34, 35]. These findings suggest that the biological pathways contributing to cachexia may differ between males and females. Future studies incorporating both sexes will therefore be important to determine whether the BMP7-dependent cachexia phenotype observed in our model is influenced by sex-specific factors.

In summary, our study identifies BMP7 as a potential master regulator of cancer-associated cachexia. BMP7 functions upstream by increasing the expression of GDF11 and GDF15, two key mediators of the cachectic phenotype. Targeting BMP7 in advanced GC holds strong translational promise, offering a potential therapeutic strategy to recover cachexia and improve patient outcomes.

## Material and Methods

### Patients and tissue samples

A total of 20 GC patients who underwent gastrectomy without preoperative treatment at Kumamoto University Hospital (Kumamoto, Japan). Immunohistochemical (IHC) staining was performed on GC specimens from eligible patients. All patients signed an informed consent form prior to participation in this study, and the study was approved by the Medical Ethics Committee of Kumamoto University (IRB approval number: 1291).

### Animal experiments

Eight- to ten-week-old male BALB/c nude mice (Clea Japan, Shizuoka, Japan) were housed in a room under stable temperature and humidity on a 12 hour-light and dark cycle. Both water and food were supplied adlibitum. All animal procedures and studies were conducted in accordance with the Institutional Animal Care and Use Committee (IACUC) at Kumamoto University (Approval number: A28-052). NUGC3, OCUM-2MD3, and MKN45 cells (5.0 × 10^6^) were implanted through intraperitoneal injection. Tumor weight and body weight were evaluated in these mice.

### Grip strength measurements

Grip Strength Meter was used to determine the maximum force displayed by the animal. The test for each mouse was repeated 3 times with a 120 s pause between each measurement. The grip strength was determined by averaging the readings from the 3 repetitions.

### Food intake measurement

Food intake was measured in singly housed OCUM-2MD3-bering mice or MKN45-bearing mice between day 14 and day 15 after tumor implantation. Pre-weighed food pellets were provided to each cage at 09:00 on day 14, and the remaining food was weighed 24 hours later at 09:00 on day 15. Daily food intake was calculated using the following formula: food intake (g/day) = amount supplied − amount remaining. Food spillage was collected from a tray placed beneath the food hopper and included in the calculation to correct for spillover.

### Cell proliferation assay

NUGC3, OCUM-2MD3, and MKN45 were seeded in 96-well plates at a density of 3.0 × 10^3^ cells/well and incubated at 37°C with 5% CO_2_. Cell proliferation was then measured with an IncuCyte S3 Livecell Analysis system (Essen BioScience). The plate was inserted into the IncuCyte instrument for real-time imaging, with four fields imaged per well every 3 hours over 4 days. Data were analyzed using the IncuCyte software, which quantified the percentage of red confluence values. All groups were tested in triplicate, and the data are presented as the means ± standard errors of the means (SEM).

### Quantitative reverse transcription (RT)-PCR

Total RNA was extracted from dissociated cells using the miRNeasy Mini Kit (Qiagen, Hilden, Germany) according to the manufacturer’s protocol. Complementary DNA (cDNA) was reverse transcribed from total RNA using SuperScript III, RNaseOUT, Recombinant Ribonuclease Inhibitor, Random Primers and Oligo(dT)12–18 Primer (Thermo Fisher Scientific). mRNA expression was quantified using SYBR Green. The reaction was performed using a LightCycler 480 System II (Roche Diagnostics). For all quantitative RT– PCR (qRT–PCR) experiments, each data point represents an independent biological replicate. All qRT–PCR data are presented as the mean ± standard error (SE). The primer sequences are listed in Table S2.

### Lentiviral transduction

shRNA lentiviral particles for transduction were purchased from Sigma; the clone IDs were as follows: shBMP7, TRCN0000058960; shGDF11, TRCN0000058952; shGDF15, TRCN0000058389; shTHNSL2, TRCN0000163285; shTNFSF13, TRCN0000059023; and sh-control, TRC1.

For lentiviral transduction, S2-VP10 cells were cultured in complete medium containing viral particles (MOI=5) with 2 mg/mL polybrene for 24 hours. Then, cells were selected in medium containing puromycin (1 μg/mL) for 3–4 days until resistant colonies could be identified. The optimal concentration of puromycin (1 μg/mL) was determined from a kill curve of medium containing puromycin at 0, 0.25, 0.5, 1.0, 2.0, 4.0, 8.0, and 10 μg/mL.

### Recombinant proteins used for *in vitro* experiments

Recombinant human BMP7 (R&D Systems) was purchased. Cells were treated with 10 ng/ml recombinant human BMP7 *in vitro* experiment.

### Western blotting analysis

Cultured cells were lysed with RIPA buffer containing a protease and phosphatase inhibitor cocktail (Thermo Fisher Scientific). The lysate was sonicated, the debris were removed by centrifugation, and the supernatant was collected as a whole-cell lysate. Protein samples were subjected to SDS-PAGE, transferred to PVDF membranes, and blotted with the primary antibodies (p-SMAD1/5 #9516S CST, SMAD1 #6944S CST, SMAD5 #9517S CST, and β-Actin #4967 CST) in Can Get Signal Solution 1 (Toyobo) at 4°C overnight. The signals were detected after an incubation with rabbit or mouse secondary antibodies in Can Get Signal Solution 2 at room temperature for 1 hour using an ECL Detection System (GE Healthcare, Little Chalfont, Buckinghamshire, UK).

### *In silico* promoter motif and ChIP-seq analysis

Human GDF11 and GDF15 promoter regions were defined as the genomic sequences spanning −2 kb to +1 kb relative to the annotated transcription start site (TSS; hg38). Candidate SMAD-binding motifs were identified using FIMO (MEME Suite) with the JASPAR SMAD5 motif (MA1557.1). Publicly available SMAD1 ChIP-seq data were obtained from ChIP-Atlas and visualized using the UCSC Genome Browser. Because comparable public SMAD5 ChIP-seq datasets were not available, publicly available SMAD1 ChIP-seq data were used for comparison with the predicted SMAD5 motif locations.

### Immunostaining and scoring methods

Paraffin-embedded sections (4 μm) obtained from GC patients were deparaffinized and soaked in distilled water. Autoclave-induced antigen retrieval was performed. Endogenous peroxidase activity was blocked using 3% hydrogen peroxide. Sections were incubated with primary antibodies against BMP7 (# PA5-11720, Thermo) overnight at 4°C. The sections were subsequently incubated with a biotin-free HRP enzyme-labeled polymer of the Envision Plus detection system (Dako, Tokyo, Japan) for 1 hour at room temperature. Positive reactions were visualized using a diaminobenzidine solution, followed by counterstaining with Mayer’s hematoxylin. All IHC staining was scored for both the intensity and percentage of cell staining. The average intensity of positively stained cells was given an intensity score from 0 to 3 (0: none, 1: weak, 2: moderate, and 3: strong expression). The average proportion of positively stained cells was estimated and given a percentage score on a scale from 1 to 6 (1 = 0-5%, 2 = 6-20%, 3 = 21-40%, 4 = 41-60%, 5 = 61-80%, and 6 = 81-100%). The two scores were multiplied to characterize 15-PGDH, CD163, CD86, PD-1 and Ki-67 expression as low (0-7) or high (8-18). The positively stained area was measured as a percentage. All scoring assessments were performed independently by two investigators.

### Analysis of The Cancer Genome Atlas and published datasets

Publicly available transcriptomic and clinical data were obtained from The Cancer Genome Atlas (TCGA) stomach adenocarcinoma (STAD) cohort. Patients with diffuse-type GC were selected based on histological annotation. Disease stage was defined according to the TNM classification, and only patients with stage II–IV disease, referred to as advanced GC, were included in the survival analysis. BMP7 mRNA expression levels were extracted from normalized RNA sequencing data. Patients were stratified into high and low BMP7 expression groups based on the median expression value. Overall survival was analyzed using the Kaplan–Meier method, and survival differences between groups were evaluated using the log-rank test.

### Statistical analysis

All experiments were performed in triplicate, and the data shown are representative of consistent results. Data are presented as the mean ± SE. The Mann–Whitney U test was used to compare continuous variables between two groups. Categorical variables were compared using the χ2 test. Correlations were evaluated by Pearson correlation coefficient analysis. Survival curves were generated using the Kaplan–Meier method, and the log-rank test was performed for survival analysis. Statistical significance was defined as p values lower than 0.05, and all data met the assumptions of the statistical test used for distribution and variance. Variance was not statistically significant in any of the results. No statistical method was used to predetermine the sample size for any of the experiments (in vitro or in vivo). All statistical analyses were performed with SPSS (Version 26, IBM Corp.) and Prism software (Version 9.0.1, GraphPad).

## Abbreviations

BAT: brown adipose tissue
BMP4: bone morphogenetic protein 4
BMP7: bone morphogenetic protein 7
GC: gastric cancer
GDF11: growth differentiation factor 11
GDF15: growth differentiation factor 15
GDNF: glial cell line-derived neurotrophic factor
GFRAL: GDNF family receptor alpha-like
OP-1: osteogenic protein-1
OS: overall survival
TCGA: The Cancer Genome Atlas
TGF-β: transforming growth factor-beta
WAT: white adipose tissue

## Conflicts of Interest

The authors declare that they have no competing interests to disclose.

## Author Contributions

Conception and design: T.Y.; data acquisition: M.Y., T. Shoda., A.N., T.Y.; manuscript writing, review, and/or revision: M.Y., T.Y., and T.I.; administrative, technical, or material support: A.Y., K.M., Y.O., and T.T.; and study supervision: YA.W., and T.I.

## Acknowledgments

TY is supported by Takeda fellowship.

This work was supported by the FOREST program of the Japan Science and Technology Agency (JST, grant no. JPMJFR200H to T. I), the P-PROMOTE program of Japan Agency for Medical Research and Development (AMED, grant no. 25ama221442h0001 to T.I.), the Japan Society for the Promotion of Science (JSPS, KAKENHI grant nos. 26K02477 to T. Y.; 24K22151, 24KK0162 and 23H02772 to T.I.), the Uehara Memorial Foundation and the Vehicle Racing Commemorative Foundation.

Illustrations were in part generated using BioRender.com.

## Data Availability

To compare cytokine expression among the GC cell lines, the **GSE22183** dataset was downloaded from the Gene Expression Omnibus.

## Notes

### Competing Interest Statement

The authors have declared no competing interest.

### Summary of Updates

Figure 4 was newly assembled by relocating the BMP7 knockdown, recombinant BMP7 rescue, and SMAD1/5 signaling data previously presented in Figure 3H-J to Figure 4A-C. New panels Figure 4D and 4E were added to show the JASPAR SMAD5 motif used for FIMO scanning and the in silico analysis of candidate SMAD-responsive regions in the GDF11 and GDF15 promoters, together with publicly available SMAD1 ChIP-seq peaks. Figure 3 was reorganized, and Figure 4 was newly assembled accordingly. The corresponding Results, Materials and Methods, Discussion, and figure legends were also revised.

