## Supplementary Information for "Identification of Bone Morphogenetic Protein 7 as a Master Regulator of Gastric Cancer-Associated Cachexia"

### Figure Legends

**Figure S1. MKN45-bearing mice exhibit significantly reduced survival compared to mice bearing other diffuse-type GC cell lines.**
A. Representative images of mice bearing MKN45, OCUM-2MD3 or NUGC3 tumors at the time of sacrifice.
B. Overall survival of mice bearing diffuse-type GC cell lines over 30 days (MKN45: n=13 mice; NUGC3: n=5 mice; OCUM-2MD3: n = 5 mice; KATOIII: n = 5 mice; OCUM12: n=5).

**Figure S2. Generation of cell lines with shRNA-mediated knockdown of BMP7, GDF11, GDF15, THNSL2, and TNFSF13.**
RT-qPCR analysis of BMP7, GDF11, GDF15, THNSL2, and TNFSF13 mRNA in MKN45 cells transfected with shRNA targeting each gene, compared to MKN45 cells transfected with control shRNA.
**p < 0.01; Mann–Whitney U test.

**Supplementary Table 1.**
**Baseline clinical and demographic characteristics of patients stratified by BMP7 expression**

| Variable | High BMP7 (n = 13) | Low BMP7 (n = 7) | p-value |
|---|---|---|---|
| Gender, n (%) | | | 1.00 (Fisher's exact) |
| Male | 10 (76.9%) | 5 (71.4%) | |
| Female | 3 (23.1%) | 2 (28.6%) | |
| Histology, n (%) | | | 0.3498 (Fisher's exact) |
| Diffuse | 5 (38.5%) | 5 (71.4%) | |
| Intestinal | 8 (61.5%) | 2 (28.6%) | |
| Depth of invasion, n (%) | | | 0.8060 ($\chi^2$ test) |
| MP | 1 (7.7%) | 1 (14.3%) | |
| SS | 8 (61.5%) | 5 (71.4%) | |
| SE | 3 (23.1%) | 1 (14.3%) | |
| SI | 1 (7.7%) | 0 (0%) | |
| Neoadjuvant therapy | Not available | Not available | — |

### Supplementary Table 2. Primer information

| Organism | Official Symbol | Alias | Forward primer | Reverse primer |
|---|---|---|---|---|
| Human | *BNP7* | | gcccaagaaccaggaagc | ggtctcggaagctgacataca |
| | *GDF11* | | accaccgagaccgtcattag | agggctgccatctgtctg |
| | *GDF15* | | cggaaacgctacgaggac | agagatacgcaggtgcaggt |
| | *THNSL2* | | ttggacacacatcccctacc | gcctatcttttgagcaatgtacc |
| | *TNFSF13* | | tcagccagtttcttgcttcc | ggtgactgaaggtgggaaag |
| | *NRP1* | | taccctgagaatgggtggac | cgtgacaaagcgcagaag |
| | *EDN1* | | gctcgtccctgatggataaa | ccatacggaacaacgtgct |
| | *BMP4* | | taccctgagaatgggtggac | cgtgacaaagcgcagaag |
| | *FAM3B* | | gaactcccttccgaaattca | tctttgggtatgcagccttc |
| | *FAM3C* | | gagggatcaatgttgccttg | aaatggtgccacatctcctc |
| | *XCL1* | | cactctccttgcacagctca | tctgagacttcactccctacacc |
| | *XCL2* | | actctccctgcacagctcag | tgagacttcactccctacacctt |
| | *TNFSF12* | | acaggatggagcacaagca | caatctggcggtcgtagc |
| | *ACTB* | β-actin | attggcaatgagcggtt | cgtggatgccacaggact |
| Mouse | *MuRF1* | | tcctgatggaaacgctatggag | attcgcagcctggaagatgt |
| | *Atrogin1* | | tcagagaggcagattcgcaa | gggtgaccccatactgctct |
| | *Cyclo* | | ggagatggcacaggaggaa | gcccgtagtgcttcagctt |

Figure S1

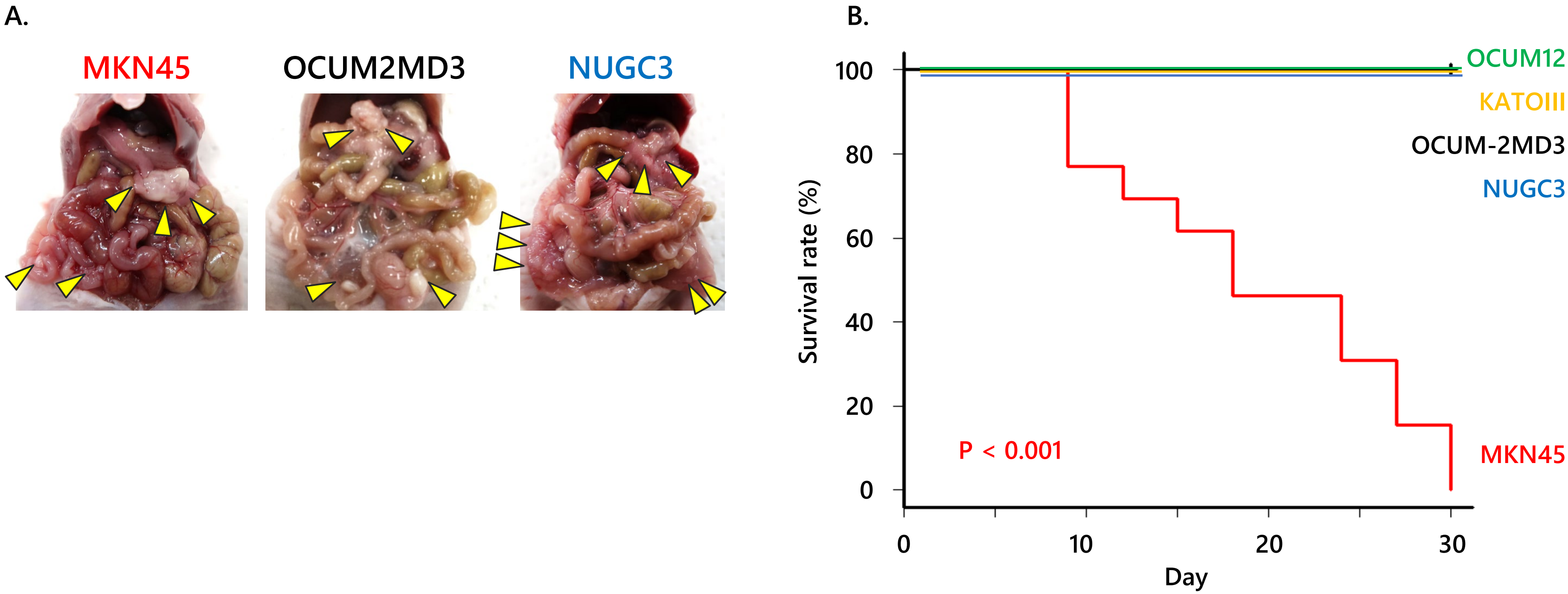


Number at risk

| | | | | | | | |
|---|---|---|---|---|---|---|---|
| MKN45 | 13 | 13 | 10 | 9 | 6 | 4 | 2 |
| OCUM12 | 5 | 5 | 5 | 5 | 5 | 5 | 5 |
| KATOIII | 5 | 5 | 5 | 5 | 5 | 5 | 5 |
| OCUM-2MD3 | 5 | 5 | 5 | 5 | 5 | 5 | 5 |
| NUGC3 | 5 | 5 | 5 | 5 | 5 | 5 | 5 |

Figure S2

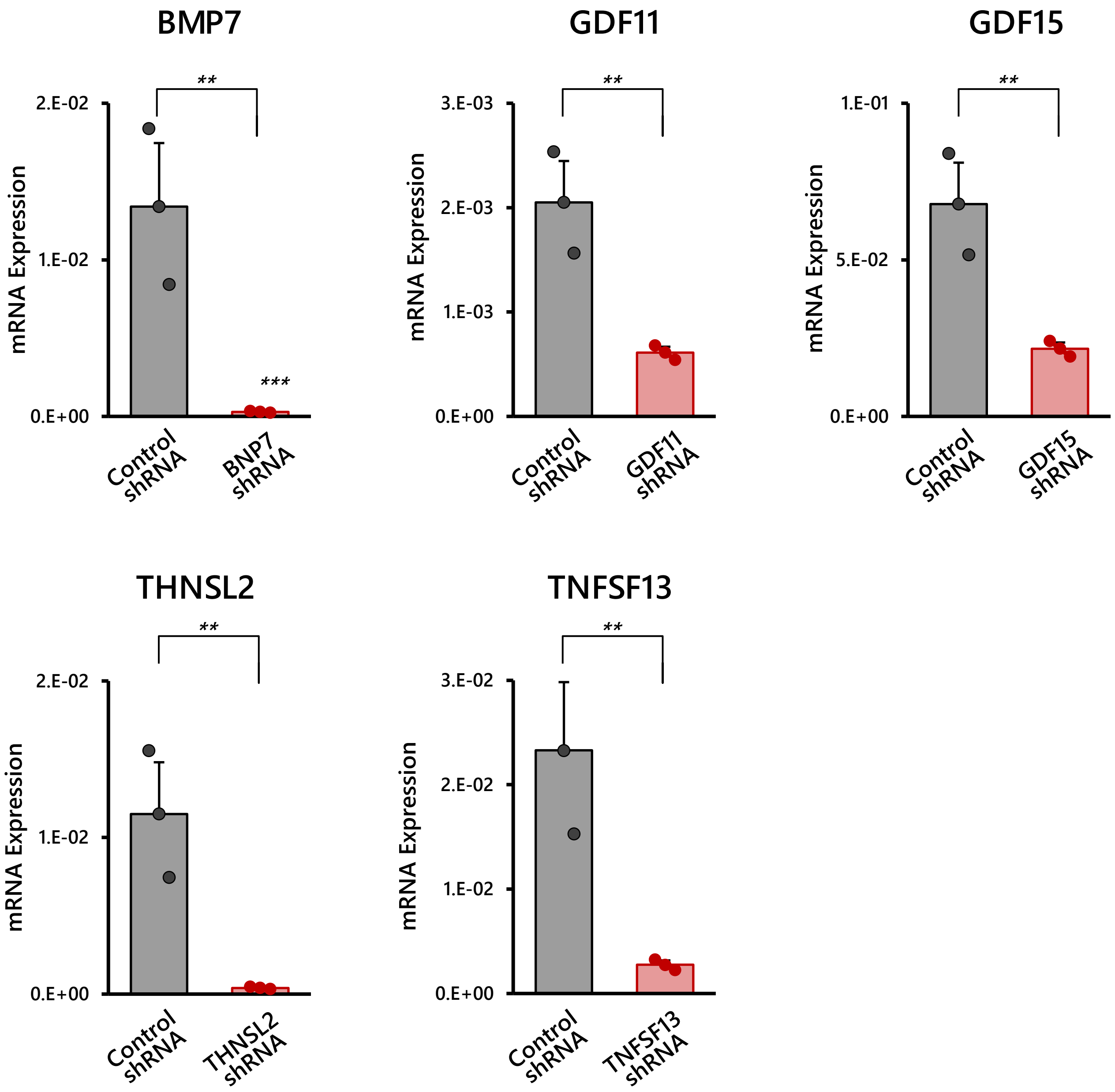

BMP7
GDF11
GDF15
THNSL2
TNFSF13
mRNA Expression
**
***
2.E-02
1.E-02
0.E+00
3.E-03
2.E-03
1.E-03
1.E-01
5.E-02
3.E-02
Control shRNA
BNP7 shRNA
GDF11 shRNA
GDF15 shRNA
THNSL2 shRNA
TNFSF13 shRNA
